# Automatic Landmark-guided Bijective Brain Image Registration by Composing Region-based Locally Diffeomorphic Warpings

**DOI:** 10.1101/2020.04.16.045609

**Authors:** Hengda He, Qolamreza R. Razlighi

## Abstract

As the size of the neuroimaging cohorts being increased to address key questions in the field of cognitive neuroscience, cognitive aging, and neurodegenerative diseases, the accuracy of the spatial normalization as an essential pre-processing step becomes extremely important in the neuroimaging processing pipeline. Existing spatial normalization methods have poor accuracy particularly when dealing with the highly convoluted human cerebral cortex and when brain morphology is severely altered (e.g. clinical and aging populations). To address this shortcoming, we propose to implement and evaluate a novel landmark-guided region-based spatial normalization technique that takes advantage of the existing surface-based human brain parcellation to automatically identify and match regional landmarks. To simplify the non-linear whole brain registration, the identified landmarks of each region and their counterparts are registered independently with large diffeomorphic (topology preserving) deformation via geodesic shooting. The regional diffeomorphic warping fields were combined by an inverse distance weighted interpolation technique to have a smooth global warping field for the whole brain. To ensure that the final warping field is diffeomorphic, we used simultaneously forward and reverse maps with certain symmetric constraints to yield bijectivity. We have evaluated our proposed method using both simulated and real (structural and functional) human brain images. Our evaluation shows that our method can enhance structural correspondence up to around 86%, a 67% improvement compared to the existing state-of-the-art method. Such improvement also increases the sensitivity and specificity of the functional imaging studies by about 17%, reducing the required number of subjects and subsequent costs. We conclude that our proposed method can effectively substitute existing substandard spatial normalization methods to deal with the demand of large cohorts and the need for investigating clinical and aging populations.

## I. Introduction

SPATIAL normalization is an essential pre-processing step in many neuroimaging studies that makes between-subjects and between-groups comparisons possible by warping each subject’s brain image onto a common or standard space. Spatial normalization is often performed by an underlying subject-to-subject or subject-to-standard image registration. Image registration, by definition, serves to make all subjects’ neuroanatomical regions correspond to a standard space and consequently to each other. Without neuroanatomical correspondences, it is challenging if not impossible, to perform any across-subjects univariate or multivariate statistical analyses (likely the most essential step in obtaining and interpreting scientific results from neuroimaging data). Yet inter-subject registration of the brain, especially the cerebral cortex, remains challenging due to its highly convoluted patterns of sulci and gyri with large inter-subject morphological variability. For example, cortical folding (e.g. sulci branches) is not consistent between subjects in many cortical regions [1]. This not only makes their registration challenging, but also increases the likelihood of false-positive findings in neuroimaging studies [2]. Better correspondence between neuroanatomical regions will improve the statistical power to detect any brain effect and will increase spatial specificity, resulting in a reduced number of required subjects and consequently the study costs [3].

The most commonly employed spatial normalization methods perform either a volume-based non-linear registration of structural images in 3D Euclidean space, or a surface-based non-linear registration of the cerebral cortex surfaces in 2D parametric surface space. For example, currently widely-used volume-based brain image registration methods include large deformation diffeomorphic metric mapping (LDDMM) [4], advanced normalization tools (ANTS) [5], Quicksilver [6], and VoxelMorph [7]. Alternatively, commonly used surface-based methods include FreeSurfer [8] and spherical demons [9]. There are also volume and surface hybrid methods available, that extend the cortex correspondence specified using surface-based registration to 3D Euclidean space. For example, Joshi *et al*. and Lepore *et al*. used harmonic mapping, whereas Postelnicu *et al*. used Navier operator of elastic diffusion for extending the surface correspondence to the volumetric space [10]-[12].

The following lists the major concerns of volume-based spatial normalization: 1) Human neuroanatomical regions are distributed throughout the highly folded surface of the cerebral cortex which often makes the Euclidean distance less reliable in segregating adjacent regions with completely distinct functionality. For instance, the posterior and anterior banks of the *Sylvian fissure* have completely distinct functionalities, but they could be seen adjacent to each other in the Euclidean space. Therefore, even a small misalignment in the Euclidean space can exert drastic consequences in matching neuroanatomical regions. 2) There is almost no difference in the intensity of the cerebral gray-matter throughout the entire cortex. This makes intensity-based similarity measures, used often as the cost function in image registration, less sensitive in distinguishing different neuroanatomical regions throughout the cortex, particularly in adjacent ones. As we have demonstrated in our simulation, even if volume-based registration were able to align the cortical folding patterns between-subjects, it would still be less likely to correspond perfectly between different cortical regions along the aligned cortical gray-matter ribbons. 3) Every non-linear image registration method relies on an underlying optimization step. Due to the complexity and large inter-subject variability in regional cortical morphology, the optimization objective function becomes nonconvex almost at all cases causing many local optima that could be detected as the final solution, without providing an accurate correspondence between the regions. Thus, even the best performing volume-based non-linear registration (ANTS) results in a poor correspondence between the cortical regions (Dice similarity coefficient (DSC) of 0.6 to 0.7 [13]). Such mismatching can attribute the location of brain activation to different regions of the standard space in group-level analysis, reducing the statistical power available to detect significant effects.

To address these shortcomings, surface-based methods were proposed to directly align brain folding patterns of gyri and sulci based on their underlying curvature instead of relying on voxel intensity. For instance, Fischl *et al*. demonstrated that compared to nonlinear volumetric methods, a surface-based method more consistently aligns brain cyto-architectonic boundaries (Brodmann areas) [14]. Optimization in surface-based methods is more efficient as it works in a 2D surface space with fewer degrees of freedom. However, all surface-based spatial normalization methods are required to project functional activation extracted from gray-matter volume to a cortical surface. This mapping process is challenging in practice and potentially problematic. For example, cortical surfaces are typically extracted from structural scans and projected onto functional image space. Due to excess geometric distortion in fast acquisition techniques, such as echo planner imaging (EPI) often used for fMRI acquisition and their low resolution, co-registration between functional and structural scans is likely to have inaccuracies that directly result in sampling non-gray-matter regions. By sampling non-gray-matter regions or regions from a neighboring gyrus/sulcus onto the cortical surface, functional activation can easily get lost and affect the results of the group-level analysis. It has been shown that when mapping functional activation from the volume to the surface, the functional signal can be diluted to neighboring gyri. This effect can be consistent across subjects and detected at the group-level, resulting in a false-positive cluster otherwise absent in volume-based spatial normalization [15]. Another shortcoming of the surface-based methods is that they cannot be applied for registering sub-cortical regions. Finally, almost all widely used brain image registration techniques whether volume- or surface-based, or a hybrid method, are based on solving the optimization problem of matching the whole brain at once, while suffering from the local minimum problem, resulting in poor correspondence between brain cortical regions.

The present study extends surface-based spatial normalization methods to the 3D Euclidean space, which is applicable to not only the cerebral cortex but also sub-cortical, cerebellar, ventricular, and other brain regions. Compared to surface-based methods, our method circumvents the projection of volumetric fMRI data onto the cortical surface. Compared to volumetric methods, instead of using voxel intensity, we directly estimate a volumetric warping field using corresponding vertices on the surface of WM/GM (WM: white matter; GW: gray matter) and GM/CSF (CSF: cerebrospinal fluid) boundaries as automatically extracted landmarks in the 3D Euclidean space, which allows us to incorporate brain anatomical information and features into the volumetric registration process. To alleviate the local minimum problem in volume, surface, or hybrid methods, we use a region-based local registration technique [16], in which each brain’s cortical/sub-cortical region is independently registered to its corresponding region. To better align the regional folding pattern within each brain region, we use surface-based spherical registration to further enhance the correspondence between different parts of the region. We then use an automatic algorithm to extract and match the correspondence of the landmarks from vertices on the WM/GM and GM/CSF boundary surfaces. For each brain cortical region, we independently estimate a landmark-guided volumetric warping field via large deformation diffeomorphisms. Instead of applying the regional warping fields individually, we propose an adaptive inverse distance weighted (IDW) interpolation for composing the locally estimated non-linear deformation fields to obtain a smooth global deformation field. Finally, we propose a symmetric registration algorithm with residual compensation similar to [17] for enforcing bijectivity into the deformation field, while matching the cerebral cortex mask and sub-cortical regions using a region-based Demons registration.

This paper is organized as follows: We first explain the detail of the proposed landmark-guided region-based spatial normalization (LG-RBSN) method in Section II. We also explain the subjects’ demographics and acquisition parameters used to acquire MRI scans in our experiments. In Section III, we first used simulated 2D images to illustrate problems associated with volume-based methods as well as demonstrating the effectiveness of our proposed method in registering these simulated special cases. We then compare our method with the state-of-the-art non-linear whole brain registration method using real human brain images. Our results show that our method achieves higher DSC than ANTS in warping the brain’s cortical regions, sub-cortical regions, and cerebral WM. In experiments with fMRI spatial normalization, our method performs better than ANTS with regard to the specificity and sensitivity of the fMRI activation at the group-level activation statistics. Finally, we include a discussion in Section IV and we conclude the paper in Section V.

## II. Method and Materials

In this section we describe the details of our novel spatial normalization method, specifically focused on accurately matching brain cortical regions. Constituting 40% of total brain mass, cortical areas are of primary interest to neuroscience when researching information-processing brain tissue [18]. Figure 1 illustrates a flowchart of different processes required for our proposed LG-RBSN method which starts with a surface reconstruction and parcellation of the cerebral cortex followed by our automatic regional landmark extraction and matching approach. Then, the landmark-guided geodesic shooting large deformation diffeomorphic registration is performed independently for each region resulting in a distinct warping field for that region. We then combine the regional warping field together using a novel interpolation technique, IDW, to give a single global warping field for the whole brain. Finally, the forward and reverse warping fields residual compensation is used to enforce bijectivity property into the global deformation field with a region-based Demons registration to keep matching of the sub-cortical regions and cerebral cortex during regularization.

**Fig. 1.**
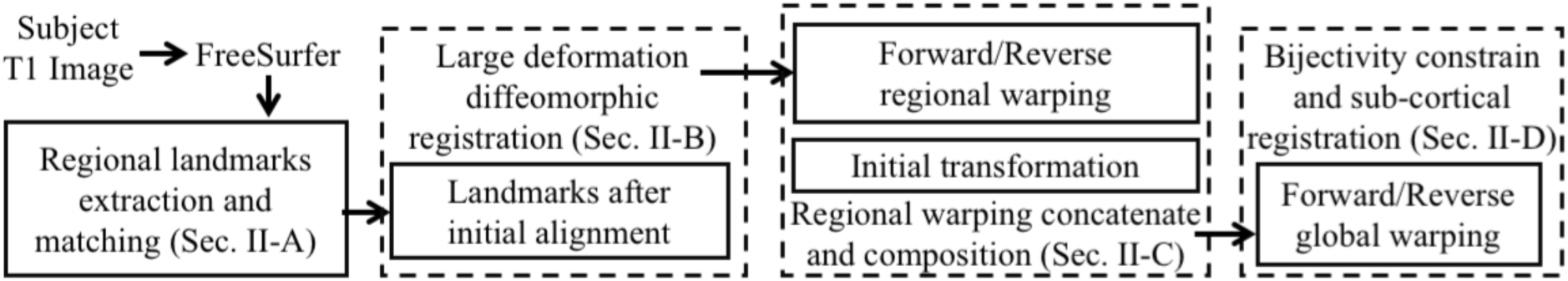
Landmark-guided region-based spatial normalization (LG-RBSN) pipeline.

### A. Automatic Regional Landmark Extraction and Matching

Landmark-based image registration can be an alternative to the volumetric registration that circumvents the use of intensity-based similarity measures to estimate a volumetric warping field. Whereas existing landmark-based image registrations generate a direct and accurate correspondence between images and do not face the local minimum problem, they require a manually identification of corresponding landmarks, which is labor intensive, subject to human error, and usually has a limited number of landmarks. For example, Anand A. Joshi *et al*. used only manually labeled sulci as landmarks to guide the registration [10], [19]. Here, we propose an automatic landmark identifying and matching procedure that accounts for approximately 2000 landmarks per region.

Our method starts with processing subjects’ structural T1-weighted images and MNI152 template using FreeSurfer pipeline, resulting in 68 cortical regions [20]. However, using FreeSurfer for initial reconstruction and delineation of the human cerebral cortex is just an arbitrary choice, other accurate surface reconstruction and delineation methods can also be used for this purpose. For each region, vertices of the WM surface (boundary between GM and WM) and pial surface (boundary between GM and CSF) triangular meshes are extracted as landmarks using the labels assigned by FreeSurfer’s cortical surface parcellation algorithm. To reduce computation for regions with large number of landmarks, the set of extracted landmarks are adaptively down-sampled with a rate varying from 10% to 60% to keep the number of landmarks around or below 2000 for each region. Down-sampled vertices are then matched back to the closest vertices on the original sphere mesh to maintain consistency between down-sampled vertices and original ones. Next, the correspondence of regional landmarks between each subject and the MNI template is established through the FreeSurfer spherical registration algorithm [8]. Specifically, the MNI vertices of each region are transformed into the subject’s spherical surface space using the spherical registration. For each projected vertex, the closest vertex of the subject’s original vertices is identified as the matching landmark in the subject’s space.

We then performed two initial linear alignments between corresponding landmarks for each region, including a global and a local transformation, independently. First, assuming that we have *m* regions *R*_*i*_ (*i* = 1, *…, m*), subject space *S*(*x*), *x* ∈ Ω_*S*_ ⊂ ℝ^3^, and MNI space *M*(*x*), *x* ∈ Ω_*M*_ ⊂ ℝ^3^, the subject’s brain structural image is registered to the MNI brain template entirely with an affine transformation *A*(*x*), and subsequently each subject’s cortical region mask is separately registered to its corresponding region in MNI space with a translation only transformation 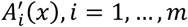. In our experiments, translation only transformations generate more appropriate linear alignments for brain cortical regions without causing any over-fitting for the subsequent regional non-linear registration. Regional landmarks are transformed using these two linear transformations for initial alignment. The framework of landmarks matching and regional registration is illustrated in Figure 2. The final regional warping field is the concatenation of two initial linear transformations and the non-linear warping field.

**Fig. 2.**
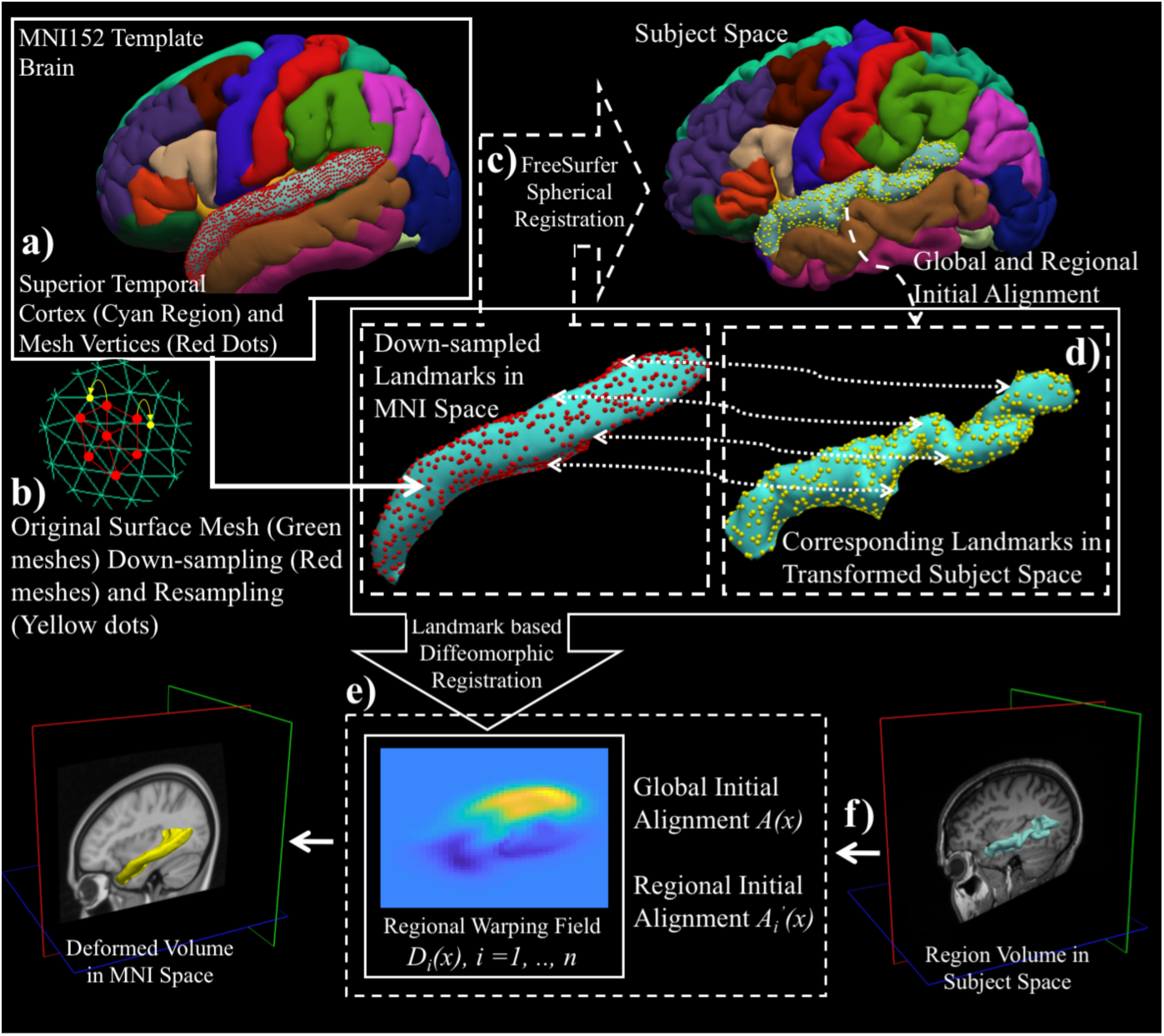
Illustrate our method for automatic landmark extraction and matching for landmark-based regional non-linear registration with example on superior temporal cortex (STC) region. In step a), for STC region (Cyan color), vertices of the GW/WM and GM/CSF boundaries triangular meshes are extracted as landmarks (GM/CSF surface vertices shown as red dots); In step b), landmarks of STC region are down-sampled by down-sampling the original dense surface mesh (green meshes) to a sparser surface mesh (red meshes) and sampled back to the original vertices (yellow dots) to keep consistency; In step c), correspondence of STC regional landmarks between the MNI template and the subject is established through spherical registration (corresponding landmarks in subject space shown as yellow dots); In step d), corresponding regional landmarks are initially aligned with linear transformations in 3D Euclidean space; In step e), a diffeomorphic non-linear landmark-based registration is used to generate regional warping field for STC region. The step f) is only showing that STC regional warping field can be used to warp the subject’s regional volume onto MNI template space.

### B. Landmark-based Large Deformation Diffeomorphic Registration via Geodesic Shooting

Traditional landmark-based non-linear image registration methods are based on smoothing spline interpolation with different radial basis functions, such as thin-plate spline [21]. Yet due to their questionable invertibility, spline-based methods often fail when dealing with the human brain’s highly convoluted topology, especially in cases that require large deformation, dense placement, or curved trajectories of landmarks [22]. To address the problem of large deformation, we use a large deformation diffeomorphism method with geodesic shooting [23] to estimate a valid warping field for each cortical region. With the optimal landmarks’ geodesic paths, both forward and reverse diffeomorphic regional transformations are estimated. In large deformation diffeomorphic registration, for the initially aligned corresponding landmarks *x*_*n*_ and *y*_*n*_ (*n* in 1, *…, N*), image warping is described as a time-varying flow quantified by a transport equation 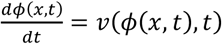. With time denoted by *t* ∈ [0,1], spatial space by *x* ∈ Ω, displacement vector field by *ϕ*(*x, t*), and velocity vector field by *v*(*x, t*). This ordinary differential equation has the solution of 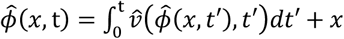, with initial condition *ϕ*(*x*, 0) = *x*. The final displacement vector field is taken as the end point of this image warping flow which is 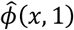.

The warping field is constrained to be diffeomorphism through a regularization penalty on the smoothness of the velocity vector field. Thus, to obtain diffeomorphic registration, the optimization objective function becomes the following:

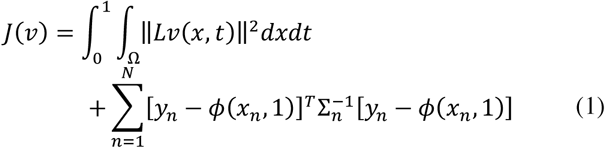

where *x*_*n*_ and *y*_*n*_ are moving and fixed landmarks, respectively, and *L* = −*a*∇^2^ + *b*∇∇ ⋅ +*c* is a linear differential operator. The second term in the objective function is the Mahalanobis distance between transformed moving landmarks and fixed landmarks, and Σ_*n*_ is the covariance matrix of landmarks, quantifying the error of inexact matching of landmarks.

Under geodesic shooting settings, the geodesic path of landmarks can be represented with an initial momentum tangent space representation at the initial time point and an initial configuration of moving landmarks. Considering the conservation law of system momentum, the objective function becomes a function of the initial momentum. The optimization problem can be solved with typical gradient descent, and the optimal landmarks geodesic path is uniquely specified using the Hamiltonian principle with the optimal initial momentum and the moving landmarks configuration. The velocity vector field is assumed as Gaussian random fields and is interpolated over the entire domain. Finally, the displacement vector field *D*_*i*_(*x*), *x* ∈ Ω_*S*_ is obtained for each region *R*_*i*_ (*i* = 1, *…, m*) with the solution of transport equation at *t* = 1. By interpolating the reverse velocity vector field from the reverse of the optimal landmarks geodesic path using the same scheme, we obtain a reverse displacement vector field 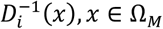. We use an available Matlab script for landmark-based diffeomorphic image registration to perform our landmark-guided non-linear regional registration [24].

### C. Inverse Distance Weighted Interpolation of Neighboring Region-based Displacement Composition

To estimate a single smooth global warping field that is applicable to all brain regions and can be applied to warp the whole brain all together at once, we propose the IDW interpolation method to combine all regional warping fields of the cortical regions. Regional deformation fields composition is illustrated in Figure 3. First, for region *i*, the global initial alignment *A*(*x*), regional initial alignment 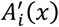, and non-linear regional deformation *D*_*i*_(*x*) are concatenated to a single regional deformation vector field 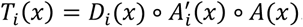. For interpolation between regions, morphology operations are performed to identify the region-to-region transition area in the brain. For example, for region *i* denoted as *R*_*i*_, all other regions *R*_*j*_ *j* = 1, *…, m, j* ≠ *i* are unioned (U_*j*_ *R*_*j*_) then dilated ((U_*j*_ *R*_*j*_) ⨁ *SE*) to intersect with the dilated region *i* (*R*_*i*_ ⨁ *SE*) for finding the transition area of the region *i* that is [*R*_*i*_ ⨁ *SE*] ∩ [(U_*j*_ *R*_*j*_) ⨁ *SE*], which leads to region *i* without transition area denoted as 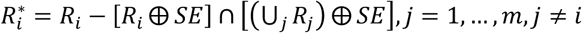, where *SE* is the structural element. Here, we use a sphere with a radius of two voxels as our *SE*. The shortest distance from spatial location *x* to 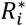, which is called *d*_*i*_(*x*), is used as the weighting factor in our IDW interpolation. The global transformation *f*(*x*) is a normalized weight of each regional displacement vector field,

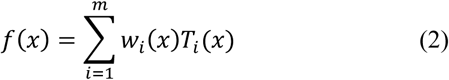

where *w*_*i*_(*x*) is the normalized weight defined as follows:

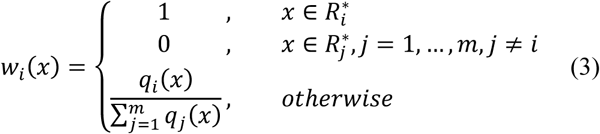

**Fig. 3.**
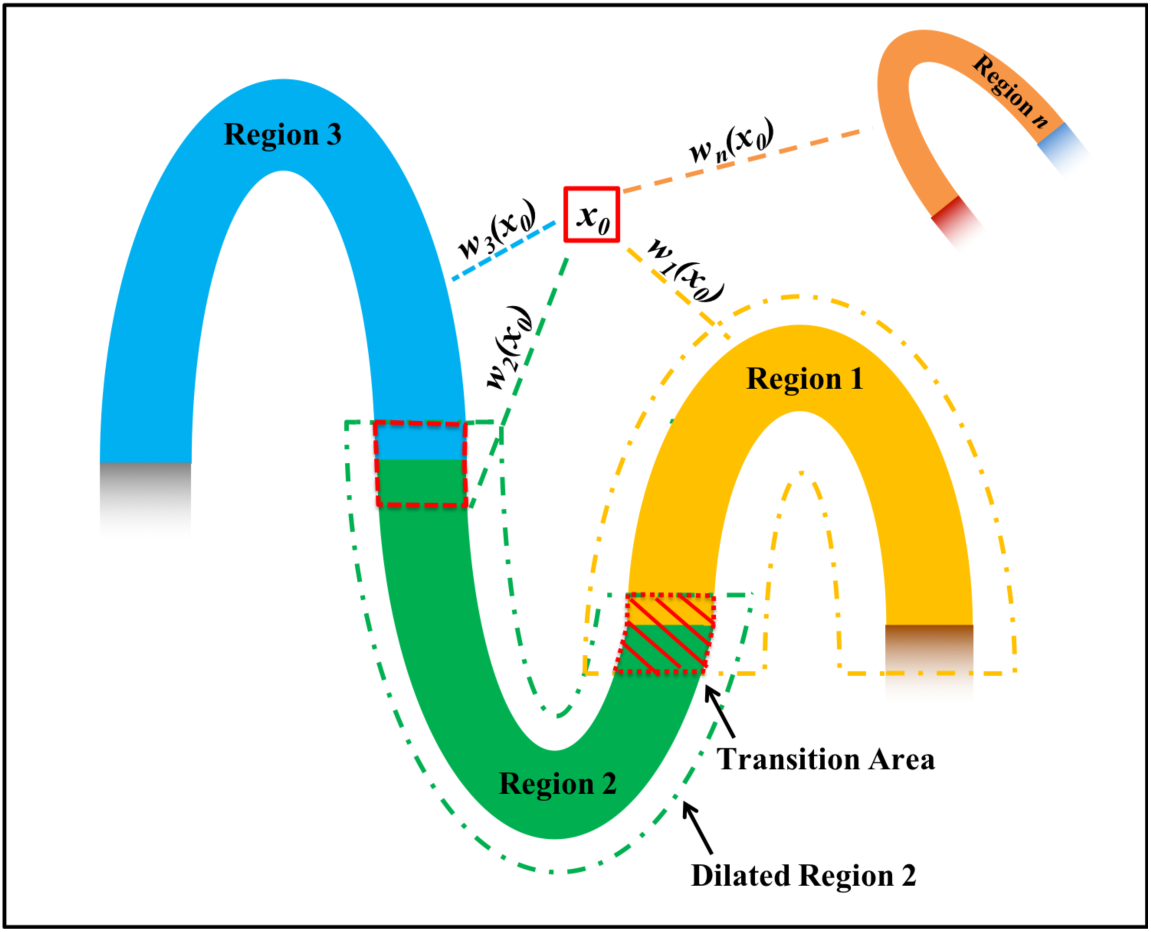
Neighboring region-based non-linear deformation fields composition using IDW interpolation. For each location in the background and region-to-region transition area, the displacement is calculated as a normalized weighted average of the displacement values in all regional warping fields at that location. The weight is based on the inverse of the closest distance between the location and the region.

Here, 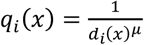 and 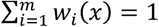. The first derivative of the interpolated global displacement vector field *f*(*x*) is continuous with *μ* > 1 [25], and in our method we set *μ* = 4. The same IDW interpolation is also applied to the reverse regional displacement vector fields 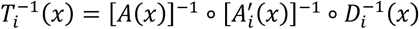 to obtain the global reverse displacement *g*(*x*). Since the IDW interpolated displacement *f*(*x*) and *g*(*x*) are no longer diffeomorphic, *f*(*x*) and *g*(*x*) are not invertible. Next, we explain our method to overcome this shortcoming in our LG-RBSN method.

### D. Bijectivity Constrain with Residual Compensation and Region-based Demons Registration

During spatial normalization the topology of a brain’s anatomical structures should be preserved between healthy normal subjects, with connected neighboring morphological structures remain connected during transformation. Topology preservation is defined as a homeomorphic map that should be continuous, bijective, and inverse continuous. Topology preservation is a desired property for the estimated deformation field to perform a valid spatial normalization, because one-to-one and bijective correspondences of location structures between one brain and another are necessary requirements for generating biologically meaningful warped brains. A simple and fast solution to embed topology preservation into the deformation field is through considering simultaneously forward and reverse maps with certain symmetric constraints. Existing topology preserving methods use different strategies to achieve this goal. For instance, Thirion used an iteration scheme to compensate for half of the residual of the reverse warped forward transformation equally to both of the forward and reverse transformations until the residual reaches identity transformation [17]. Ashburner *et al*. used a Bayesian framework with a symmetric prior, so that the probability distribution of the forward and reverse deformations is identical [26]. Inverse consistency constraint was proposed in [27], which used a symmetric cost function. Avants *et al*. proposed a symmetric diffeomorphic registration model where forward and reverse deformations meet at the middle of the registration [5], whereas Kuang used a cycle-consistent design in a deep-learning network to learn forward and reverse deformations concurrently [28].

In our method, the deformation field for each region is diffeomorphic with strict bijective constraints, but the composited deformation using IDW interpolation will no longer guarantee bijectivity. For example, moving local gyrus images needed to be cut to prevent overlap during warping [29]. To address invertibility a poly-affine method was proposed that composites local velocity vector fields associated with each local affine transformation, rather than compositing the displacement [30]. However, this velocity vector field parametrization of the displacement is difficult for non-linear transformation in our case. Alternatively, our method uses a residual compensation method to impose bijective property into the existing non-bijective global warping fields *f*(*x*) and *g*(*x*), while using a region-based Demons registration to match sub-cortical regions and the cerebral cortex mask. This prevents the mismatching of brain structures during residual compensation.

Given subject space *S*(*x*), *x* ∈ Ω_*S*_ ⊂ ℝ^3^ and MNI space *M*(*x*), *x* ∈ Ω_*M*_ ⊂ ℝ^3^, we use a residual compensation scheme to enforce bijectivity into both the direct forward displacement vector field *u*_*SM*_(*x*) = *f*(*x*), *x* ∈ Ω_0_ from *S* to *M* and the reverse displacement vector field *u*_50_(*x*) = *g*(*x*), *x* ∈ Ω_*M*_ from *M* to *S*. At the same time, we use a region-based Demons registration method to match the brain’s sub-cortical regions (*SubR*^*i*^, *i* = 1, *…, n*) and cerebral cortex mask (*CC*). We have initial direct deformation 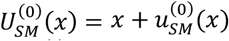 and initial reverse deformation 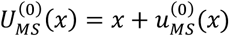. At each iteration *t*, we update both 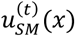 and 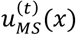 according to following steps:

i. Compute residual deformation

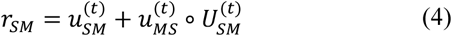

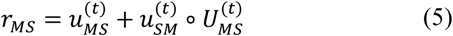
ii. Compute Demons velocity for matching the brain’s cerebral cortex mask

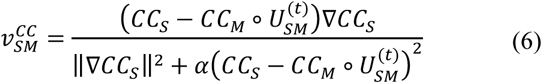

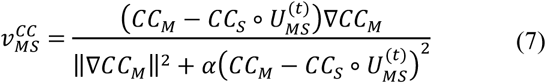
iii. Compute Demons velocity for matching the brain’s sub-cortical regions

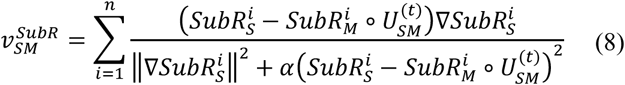

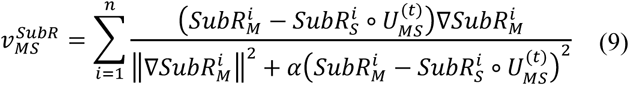
iv. Update displacement field

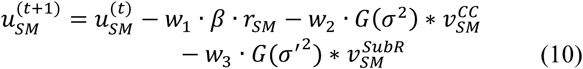

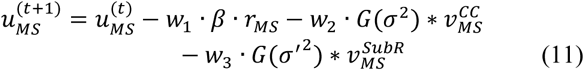

where, *w*_1_, *w*_2_, and *w*_3_ are normalized weight with *w*_1_ + *w*_2_ + *w*_3_ = 1, *G*(σ^2^) is a Gaussian kernel with variance σ^2^. We will show in the experiment section that by using this approach we can significantly reduce the number of non-positive Jacobian voxels after IDW interpolation.

### E. Subjects and Data Acquisitions

42 subjects (27/15 young/older, age (mean ± std) = 25.11 ± 3.24/66.93 ± 3.71 years) were scanned using a Siemens Prisma 3-Tesla MR scanner. T1-weighted images were acquired using a magnetization-prepared rapid gradient-echo (MPRAGE) (TR = 2300 ms; TE = 2.32 ms; flip angle = 8°, voxel size = 1 mm × 1 mm × 1 mm; matrix size = 256 × 256, and 192 slices without gap). Task-based functional MRI were acquired using a T2*-weighted multiband gradient-echo EPI (TR = 1 s; TE = 30 ms; flip angle= 62°, 64 slices without gap; slice thickness = 2 mm; 480 volumes; voxel size 2 mm × 2 mm × 2 mm, multiband factor=4) pulse sequence. Another fMRI scan was acquired in the opposite phase encoding direction, which was used in this work solely for geometric distortion correction (GDC).

We employed an event-related fMRI experimental task design. The task consisted of two ongoing stimuli: 1) A maximum contrast flashing checkerboard (i.e., visual stimulus) presented on either the right or left side of the screen, and 2) An alternating tone (i.e., auditory stimulus) paradigm played on either the right or left ear. The two sensory stimuli were presented with random onsets and durations (uniform distribution, range = 1.0 - 5.0 sec). Overlaps between visual and audio stimuli were allowed, however temporal overlapping of the bilateral presentation in the same modality were prohibited.

The data were collected in 2 runs; in the first run, subjects were instructed to attend to only one sensory stimulus (i.e., either visual or tonal) while ignoring the other. In the second run, they were instructed to attend the other sensory stimulus. Each scan consisted of 120 events: 60 events for visual and 60 for auditory stimulus. For each modality, 30 events on the right and 30 events on the left side spaced at inter-stimulus-intervals in the range of 1 - 17 sec was drawn from a uniform distribution. Control for attention was achieved by asking the subjects to press a button twice with their right/left index finger (depending on the lateralization of the attended stimulus) as soon as the attended stimulus terminated. These responses were recorded during the entire scans. Throughout the experiment, subjects were required to maintain their gaze on a minuscule fixation spot in the center of the screen, and were given feedback on any incorrect or out-of-time responses by changing the color of the fixation spot from green to red. Eye fixation was monitored by recording the eye postilion at all times using an eye-tracking system. Subjects were trained multiple times outside of the scanner to learn and perform the task properly. All subjects learned the task correctly.

## III. Experiments and Results

In this section, we evaluate the performance of the proposed LG-RBSN technique using simulated and real data. We first generated simulated 2D images of folded ribbons that resembles the folding patterns of the human cerebral cortex. Our main goal in this simulation was to show how state-of-the-art normalization methods could fail in registering such simple 2D scenarios. We then extended our validation to experiments with real MRI and fMRI data targeting both primary visual and audio regions. Our results were compared to a state-of-the-art non-linear whole brain registration method (ANTS) that is currently considered the top performing non-linear registration algorithm [13].

### A. Simulation of Cortical Gyrus Registration

To illustrate the problems associated with volumetric whole-brain registrations and to demonstrate the effectiveness of our proposed method to overcome those problems, we simulate a single cortical gyrus registration experiment in 2D. To best simulate the real task of cortical regions registration we assumed that both moving and fixed gyri have a similar shape and width (4 pixels corresponding to 4 mm in most of the currently acquired T1-weighted MRI scans), but are comprised of 3 regions of different lengths across the gyrus, as shown in the first column of Figure 4 where the 3 cortical regions are color-coded for both images. The two rows in Figure 4 illustrate two simulated experiments with different initial positions: (a) with an aligned initial position, and (b) with a mis-aligned initial position. Please note that even with initial affine registration of the whole brain, many cortical features (sulci and gyri) could still remain completely misaligned. Therefore, the simulated initial positions of alignment and mis-alignment between the moving and fixed images are very common in most spatial normalization or brain image registration techniques. We used regional landmarks distributed at two sides of the ribbons in each region, as shown in the second column of Figure 4, to estimate a global bijective warping field using our method LG-RBSN. We also applied a modified ANTS non-linear registration pipeline from the half-C to full-C registration experiment to match the moving and fixed images [31].

**Fig. 4.**
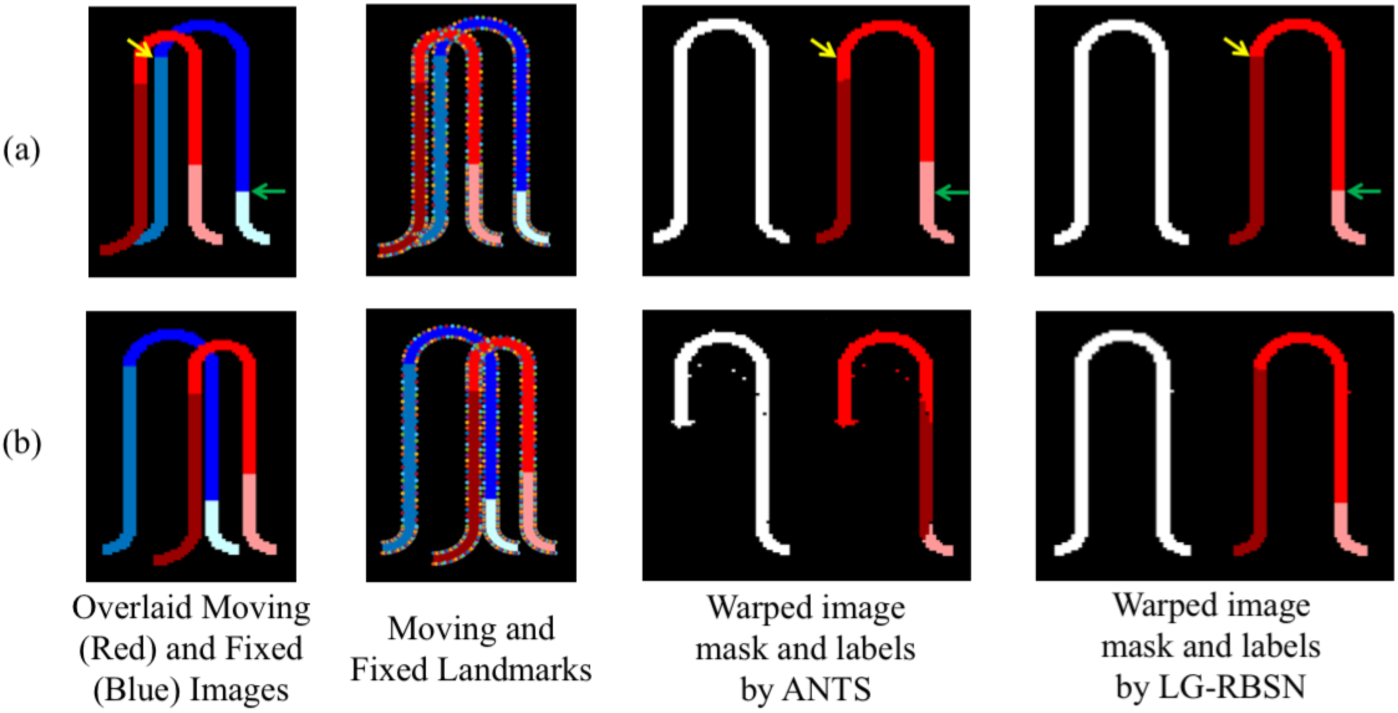
Comparison of ANTS and LG-RBSN registration results in cortical gyrus registration simulations with cases of moving and fixed images (a) aligned initially and (b) mis-aligned initially. In experiment (a), ANTS matched the whole gyrus mask perfectly but failed to accurately align the underlying 3 regions (yellow and green arrows mark the same location across images). In experiment (b), ANTS fell into a local minimum and failed to match even the binary mask of the entire gyrus. LG-RBSN matched the whole structure mask and regions perfectly in both experiments.

In experiment (a), both LG-RBSN and ANTS resulted in a comparable overlap between the entire gyrus structure (ANTS: 99.11% versus LG-RBSN: 99.86%), shown as a binary mask in the third and last columns of Figure 4. However, such high correspondence for the entire gyrus, did not hold for the 3 comprising regions and the average regional DSC dropped from 99.91% for LG-RBSN to 86.39% for ANTS. This is because intensity-based volumetric registrations, such as ANTS, cannot discriminate adjacent cortical regions. Therefore, the underlying 3 regions will not necessarily be registered, as shown in the third column of Figure 4. Alternatively, LG-RBSN almost perfectly aligns the underlying regions, as shown in the last column of Figure 4.

In experiment (b), unlike LG-RBSN which generated a perfect correspondence, ANTS failed to generate an acceptable overlap even on the binary mask of the entire gyrus (ANTS: 82.49% versus LG-RBSN: 99.66%), as seen in third and last columns of Figure 4. As mentioned in the introduction, the ANTS failure is due to its vulnerability to the local minimum during optimization. The global affine initial alignment failed to provide a sufficient initial alignment of cortical regions, which is often the case in any non-linear registration problem. Consequently, the underlying 3 regions drastically failed to correspond when using ANTS methods (ANTS: 42.60% versus LG-RBSN: 99.66%), as seen in the third and last columns of Figure 4. These results highlight the importance of detecting a true optimum point in any non-linear registration method.

In addition to the computed DSC measures for all aforementioned experiments, Table I also lists the number of voxels with non-positive Jacobians in each registration method. Jacobian matrix is used to evaluate the diffeomorphic property of the warping field, as the local deformation is invertible and preserves the topology only at locations with positive Jacobian determinant. Whereas ANTS produces 217 pixels with non-positive Jacobian determinant in experiment (b), all pixels show positive Jacobian determinant when LG-RBSN is being used for registration, emphasizing the performance of a residual compensation method in our LG-RBSN.

**TABLE 1.** DSC between warped and fixed images of using ANTS and LG-RBSN in the simulation cases of (a) aligned and (b) mis-aligned initially

| Experiments | DSC between warped moving and fixed binary mask of all regions |  | Average DSC between warped moving and fixed image labels |  | Number of non-positive Jacobian pixels |  |
| --- | --- | --- | --- | --- | --- | --- |
|  | ANTS | LG-RBSN | ANTS | LG-RBSN | ANTS | LG-RBSN |
| (a) | 99.11% | 99.86% | 86.39% | 99.91% | 0 | 0 |
| (b) | 82.49% | 99.66% | 42.60% | 99.66% | 217 | 0 |

### B. Evaluation Using Human Brain Structural Images

Using our LG-RBSN method we estimated a global registration warping field between each subject, described in Section II-E, and the MNI152 template utilizing regional WM and pial surfaces vertices to extract corresponding landmarks. For comparison, we also used ANTS (Deformation model: SyN; Similarity: normalized mutual information; Regularization: Gaussian smoothing) to perform the same registration. For qualitative evaluation of our method and ANTS, the two obtained global warping fields were applied to each subject’s T1-weighted structural brain image, with results shown in Figure 5 for three selected subjects. LG-RBSN showed a clear improvement in aligning brain cortical regions as highlighted by red dotted circles in Figure 5. The arrows in Figure 5 show how a sulcus can be generated by ANTS where the target image does not have such a structure (Subject 1), and how ANTS mismatched a gyrus of the subject to a sulcus in MNI152. Our method on the other hand properly matched the corresponding sulcus of the subject to the sulcus in MNI152 (Subject 2). This happened because ANTS performs optimization in 3D Euclidean space to find correspondence of the brain. A small shift in the volume space can mismatch two functionally distinct locations of the brain, whereas our method uses spherical registration to find the correspondence that directly matches brain folding patterns.

**Fig. 5.**
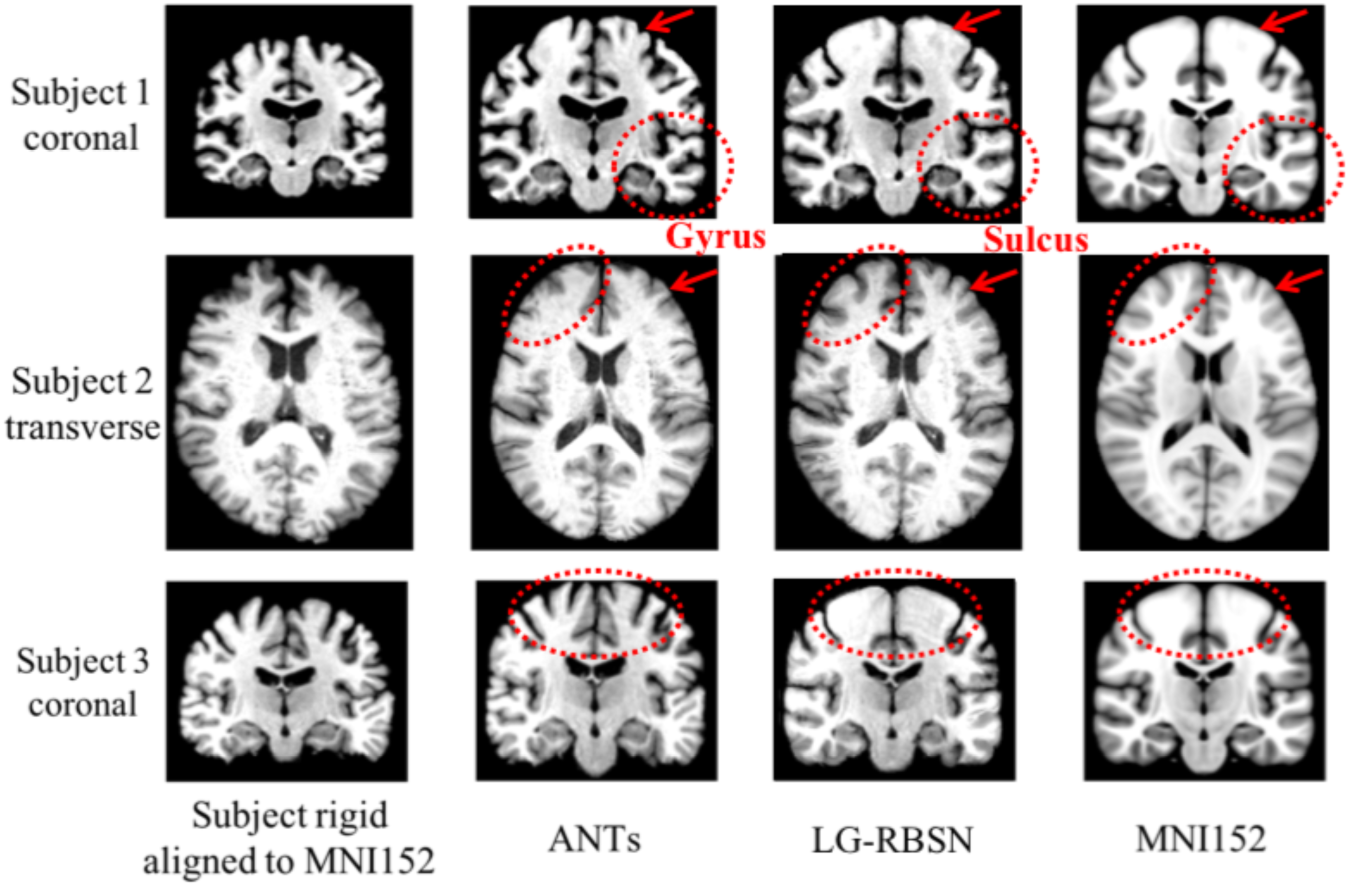
Spatial normalized brain qualification evaluation comparison between LG-RBSN and ANTS. LG-RBSN showed clearly better performance compared to ANTS in red dotted circles highlighted areas. In subject 2, ANTS mismatched a Gyrus of the subject’s cerebral cortex to a Sulcus in MNI152 space, whereas LG-RBSN matched the corresponding Sulci properly.

For quantitative evaluation, we used the global warping fields obtained above to warp each subject’s FreeSurfer delineated regions, described in Section II, onto MNI152 space (cortical regions include banks of superior temporal sulcus, caudal anterior cingulate, caudal middle frontal, corpus callosum, cuneus, entorhinal, fusiform, inferior parietal, inferior temporal, isthmus cingulate, lateral occipital, lateral orbitofrontal, lingual, medial orbitofrontal, middle temporal, parahippocampal, paracentral, pars opercularis, pars orbitalis, pars triangularis, pericalcarine, postcentral, posterior cingulate, precentral, precuneus, rostral anterior cingulate, rostral middle frontal, superior frontal, superior parietal, superior temporal, supramarginal, frontal pole, temporal pole, transverse temporal and insula). As done previously [6], [7], [13], we have evaluated our registration accuracy using the DSC between the regional binary masks of the warped and corresponding target FreeSurfer regions. Figure 6 illustrates the distribution of the DSC between corresponding regions for all cortical regions in FreeSurfer using boxplot for both LG-RBSN (in blue) and ANTS (in red) categorized in five lobar brain segments. As shown in Figure 6, LG-RBSN significantly outperforms ANTS in all cortical regions reaching to highest DSC in the insula region with (DSC = 0.9185 ± 0.0116 (mean ± std), improved by 28.35%; t = 25.34; p = 6.02e-59) and the lowest DSC in the pericalcarine region (DSC = 0.7737 ± 0.0379 (mean ± std), improved by 140.88%; t = 42.16; p = 1.36e-90). Comparing the improvement in lobar segments of the brain showed that the highest improvement was achieved in the occipital lobe (DSC improved by 108.36%; t = 53.37; p = 1.44e-165) with the least improvement in the cingulate lobe (DSC improved by 59.41%; t = 50.39; p = 3.78e-158). As shown in Figure 7, in total, using LG-RBSN has substantially improved the correspondence between cortical regions (DSC = 0.8558 ± 0.0444 (mean ± std)), and is significantly higher (DSC improved by 67.31%; t = 142.30 p = 0) than the results obtained by ANTS (DSC = 0.5115 ± 0.1214; mean ± std).

**Fig. 6.**
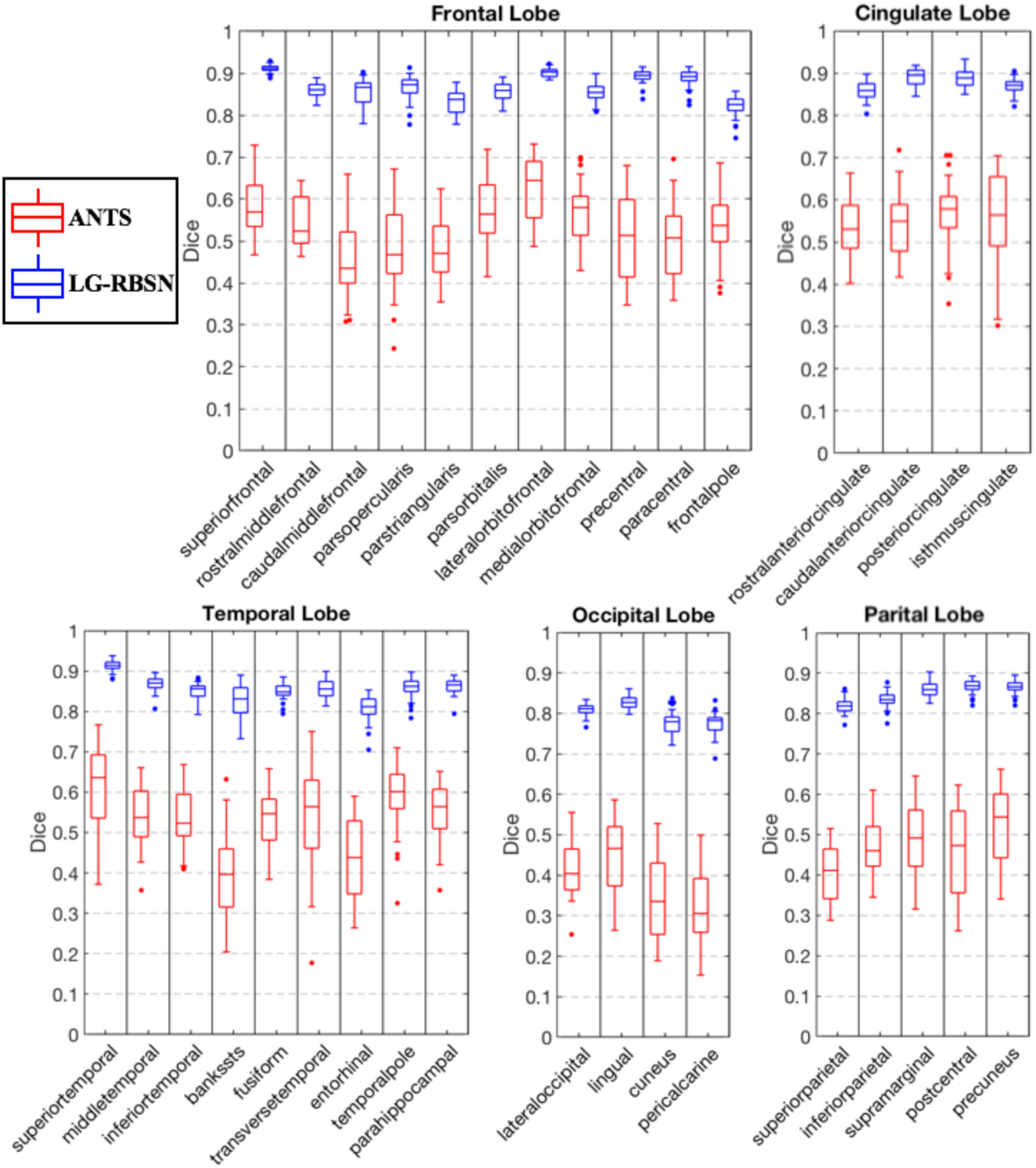
Cortical regional DSC comparison between LG-RBSN and ANTS in different brain lobes. LG-RBSN showed significantly higher DSC in matching brain cortical regions than ANTS. And LG-RBSN is more robust working with both young and older subjects compared to ANTS, as LG-RBSN showed less variance of DSC in matching brain cortical regions.

**Fig. 7.**
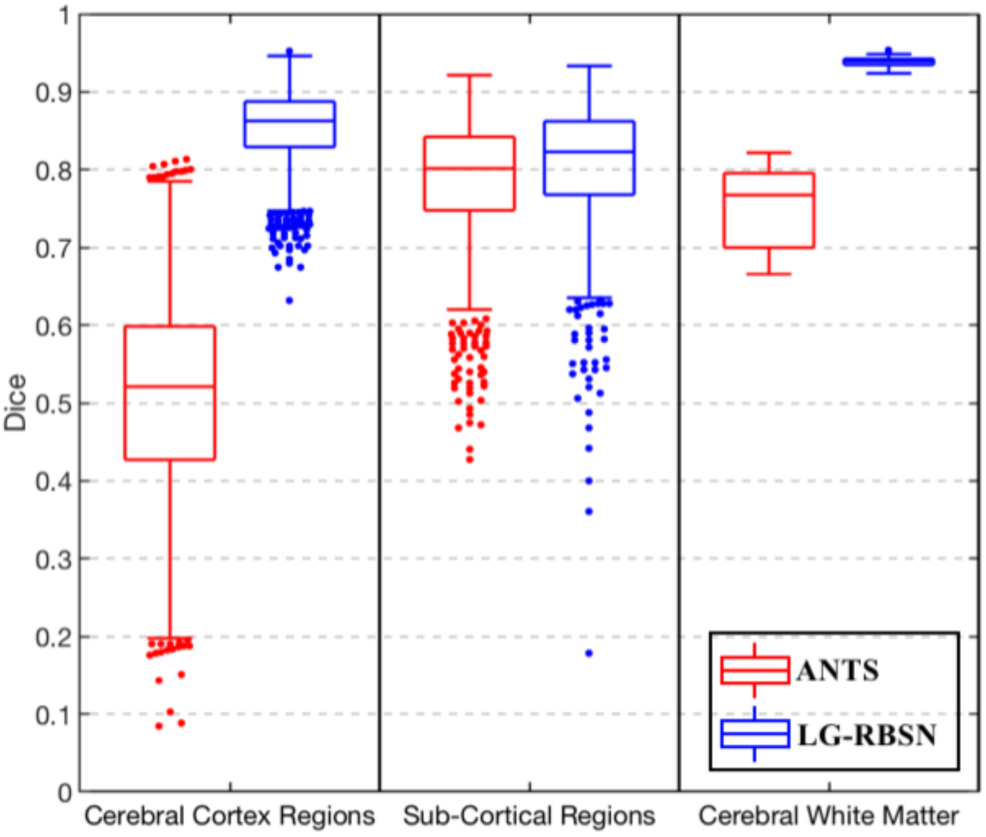
DSC comparison between LG-RBSN and ANTS. LG-RBSN showed significantly higher DSC in matching brain cortical regions, sub-cortical regions, and cerebral WM than ANTS. And LG-RBSN is more robust working with both young and older subjects compared to ANTS, as LG-RBSN showed less variance of DSC in matching brain cortical regions and cerebral WM.

Next, we evaluated the effectiveness of the LG-RBSN for aligning the sub-cortical regions and cerebral WM in comparison with the results obtained by using ANTS (segmented with FreeSurfer, sub-cortical regions include lateral ventricle, ventral DC, cerebellum white matter, cerebellum cortex, thalamus, caudate, putamen, pallidum, hippocampus, amygdala, third ventricle and brain stem). Figure 7 shows the results of this evaluation along with the evaluation of all cortical regions obtained above. As shown in Figure 7, our method significantly (t = 5.53; p = 3.71e-08) outperformed ANTS in matching sub-cortical regions with a higher DSC values (0.8056 ± 0.0823; mean ± std) compared to DSC values when using ANTS (0.7840 ± 0.0854; mean ± std). Furthermore, our method showed significantly improved DSC (0.9387 ± 0.0054; mean ± std) of matching cerebral WM mask (DSC improved by 25.55%; t = 34.86; p = 2.83e-78), compared to the DSC of using ANTS (0.7477 ± 0.0500; mean ± std). ANTS’s poor performance in warping cerebral cortex regions compared to reported DSC in the existing literature may be due to the use of the MNI152 template instead of a real subject as the target space, evaluation with small cortical regions overlap and the vulnerability of regions in temporal lobe and frontal lobe to age-rated brain atrophy and deformation in our study (about one-third of our subjects are healthy older adults).

Akin to the simulation section, the number of non-positive Jacobian voxels is used to quantify the bijectivity property of the warping field. To validate the bijectivity of the global warping field obtained with our method and ANTS, we calculated the number of non-positive Jacobian voxels along with iterations. Results are shown in Figure 8. The final warping fields estimated by using our method have number of non-positive Jacobian voxels 1338.1 ± 404.3 (mean ± std) for the forward warping field and 1318.3 ± (mean ± std) for the backward warping field, compared to ANTS with 1483.2 ± 2121.1 (mean ± std) for the forward warping field and 1473.9 ± 2239.1 (mean ± std) for the backward warping field. There is no significant difference of the number of non-positive Jacobian voxels between ANTS and LG-RBSN for both forward (t = 0.4354; p = 0.6644) and backward (t = 0.4433; p = 0.6587) warping fields.

**Fig. 8.**
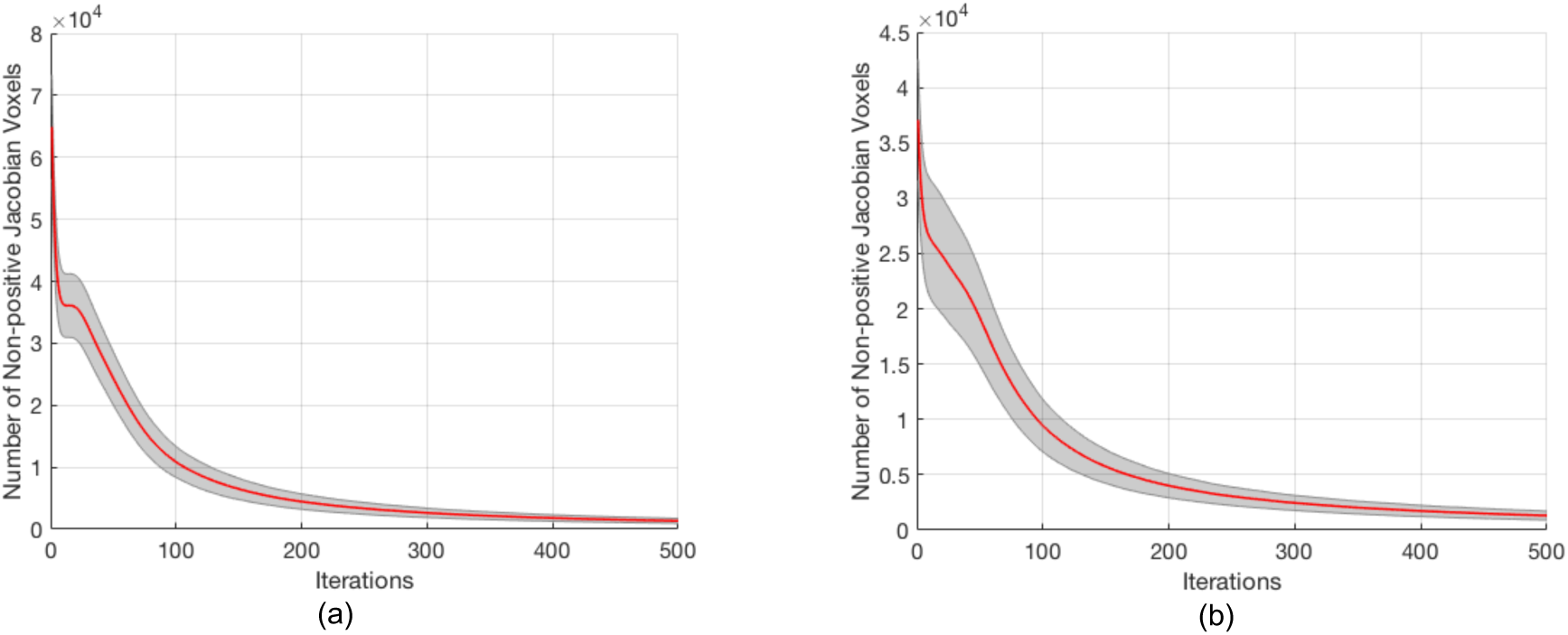
Number of non-positive Jacobian voxels decreases along bijectivity constrain iterations for (a) forward (subject to MNI152) (b) backward (MNI152 to subject) warping field. The red curve and the grey region represent the mean and the standard deviation.

### C. Evaluation Using Human Brain Functional Images

Whereas we have shown in the previous section that LG-RBSN significantly improves the regional correspondence between warped and reference images, that does not necessarily imply that the enhancement will be directly transformed to functional images. As we have shown previously in a preliminary study [16], the improvement in the structural overlap had significantly increased the statistics of the group-level brain activations in primary visual cortex, but it did not generalize to the group-level activations from the primary auditory cortex. By enhancing the LG-RBSN method in this work, we aim to overcome this problem. We again applied our method to the statistical parametric maps obtained from both an auditory and a visual fMRI experiment to generate group-level activation maps.

We have compared the obtained activation maps to those generated using ANTS, and to the anatomy of the primary visual and auditory cortices, where we expect to detect the true-positive activations. The preprocessing pipeline for the task-based fMRI data is illustrated in Figure 9. Briefly, slice timing correction is applied to the raw fMRI timeseries to account for the difference in the acquisition delay between slices [32]. At the same time, motion parameters are estimated on raw fMRI scans using rigid-body registrations performed on all the volumes in reference to the first volume. Additionally, the first volumes are extracted from another fMRI scans with opposite phase encoding directions to estimate the geometric distortion correction field using a susceptibility-induced distortions correction technique called topup [33] provided in FSL software package [34]. Then, the estimated motion parameters and geometric distortion field are combined and applied to the slice timing corrected fMRI time-series to get the distortion and motion-corrected fMRI time-series. First level general linear modeling is performed independently on each voxel using multiple regression with four variables of interest (stimuli timing convolved with canonical Double-Gamma hemodynamic response function) resulting to 4 different statistical parametric maps which will be warped onto a standard space to be able to perform group-level statistical analysis.

**Fig. 9.**
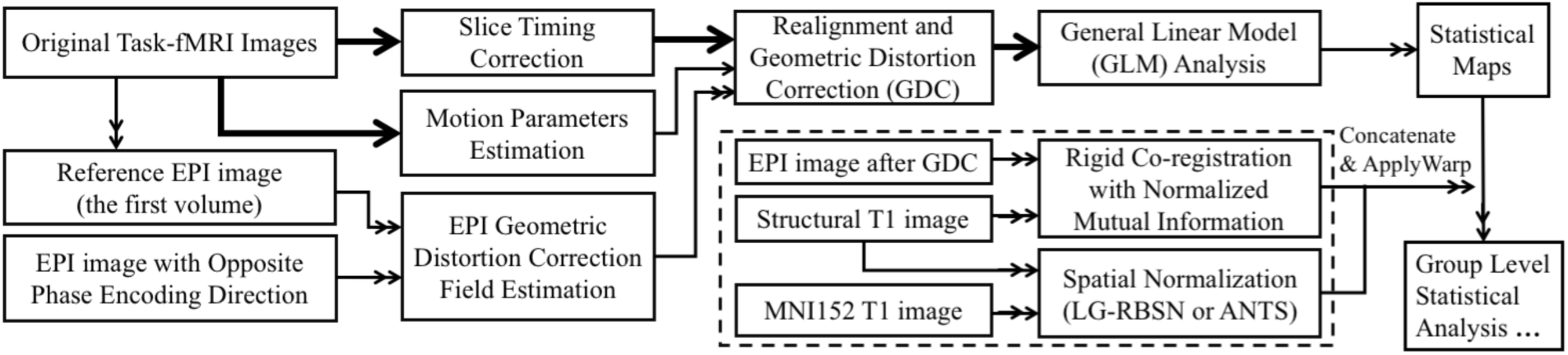
The processing pipeline for task-based fMRI data. The thick arrows shown the transfer of 4D fMRI data, the double thin arrow shows the transfer of the 3D data, and the thin arrow shows the transfer of the parameters.

Each subject’s global registration warping field estimated in the previous experiment was concatenated with the within-subject functional to structural rigid-body transformation, and used to project brain auditory/visual activation statistical maps of that individual into MNI152 space for group-level analysis. Group-level analysis was done by simple regression where a voxel was deemed active if its averaged point-estimates were significantly different from zero.

We first compared the actual statistics of the group-level analysis results when LG-RBSN was used as the spatial normalization versus the results obtained with using ANTS. The statistics are quantified as the point estimates (β values) of the 100 activated voxels with highest z-statistics in the brain’s left occipital lobe for stimulating right visual hemifield and vice versa. We have combined four FreeSurfer regions (lateral occipital, lingual, cuneus, and pericalcarine gyrus) to obtain the binary mask of the occipital lobe. Figure 10 illustrates the distribution of the point-estimates in the top 100 activated voxels using boxplots (Left: for stimulating left visual hemifield; Right: for stimulating right visual hemifield). As seen in Figure 10, the mean point estimates of group-level brain activation using our method (Left: β = 578.51 ± 33.19, Right: β = 637.89 ± 55.73) is significantly higher (Left: t = 8.64 p = 1.92e-15, Right: t = 26.94 p = 0) than that obtained by ANTS (Left: β = 534.39 ± 38.83, Right: β = 473.43 ± 24.94). These results indicate that the magnitude of the fMRI signal will significantly increase if we use a more accurate spatial normalization technique.

**Fig. 10.**
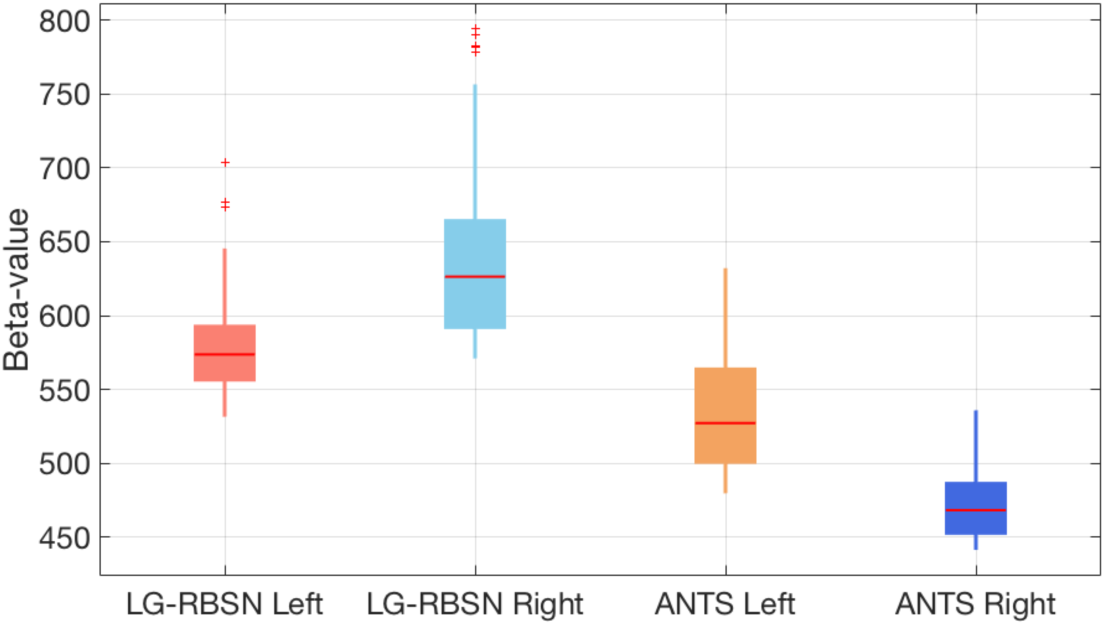
Distribution of the point estimates for visual task-based fMRI group level visual activation in brain contralateral occipital lobe using different spatial normalization methods (Left: for stimulating left visual hemifield; Right: for stimulating right visual hemifield).

For auditory stimulation we used combination of two FreeSurfer regions (transverse temporal gyrus and superior temporal gyrus) to generate a binary mask for the primary auditory cortex. Figure 11 illustrates the distribution of the point estimates from the 100 voxels with highest significance level within the binary mask using boxplots and also shows the comparison between the point estimates obtained by our LG-RBSN method and the ones obtained by using ANTS (Left: for stimulating left ear; Right: for stimulating right ear). As seen in this figure, the mean point estimates of the group level brain activation using our method (Left: β = 359.74 ± 24.32, Right: β = 304.70 ± 17.21) is significantly higher (Left: t = 17.00, p = 1.45e-40; Right t = 8.19, p = 3.16e-14) than that the one obtained by ANTS (Left: β = 308.20 ± 18.09, Right: β = 283.84 ± 18.77). These results indicate that the performance of the LG-RBSN supersede our originally implemented RBSN [16], where improvement in the regional correspondence did not transform to improvement in the group-level activation statistics.

**Fig. 11.**
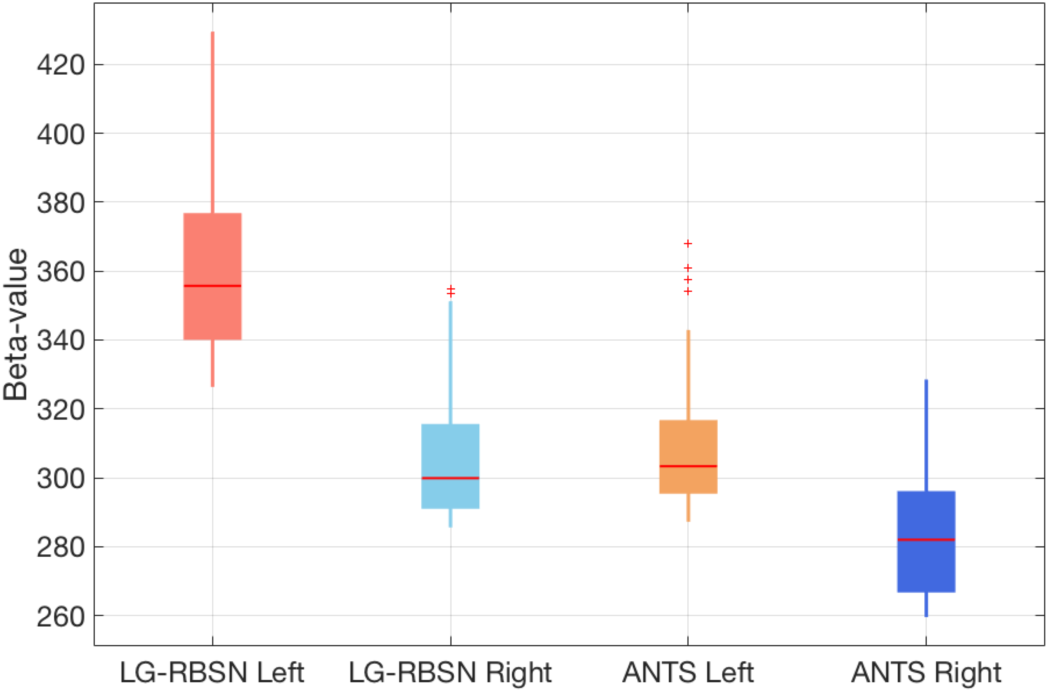
Beta values of tonal task fMRI group level auditory activation in contralateral transverse temporal cortex and superior temporal cortex using different spatial normalization methods (Left: for stimulating left ear; Right: for stimulating right ear).

Improving the group-level statistics in our fMRI experiments shows that the proposed spatial normalization technique increases the statistical power to detect smaller effects that may not be detectable with the conventional methods. Still, the method does not guarantee that the false-positive rate will not increase as well, which is often the case in many advanced developments for fMRI processing. To address this problem, we use receiver operating characteristic (ROC) curves which associate the cost of improvement in the true-positive rate (sensitivity) to the false-positive rate (1-specificity) at different threshold levels for detecting an effect. However, using ROC curve for evaluating any fMRI experiment is a challenging task due to the lack of gold standard measurement. In this paper, we use the masks of the primary visual cortex (lateral occipital) and auditory cortex (transverse temporal gyrus), given by FreeSurfer, as the regions that we expect to see activated voxels in and any detection of activated voxels outside these masks can be considered as false-positive. Therefore, true-positive rate is the number of activated voxels divided by the total number of voxels inside the region, and false-positive rate is the number of activated voxels in the vicinity of the regional masks (obtained by dilating the same regional masks) divided by the total number of voxels in the dilated regions. By changing the threshold for significance (t-statistics ranging from 0 to 15), we plot the curve illustrating the association between these two rates. The area under the curve (AUC) of the ROC is often used as the main performance metric for quantitative comparison.

Figure 12 illustrates the ROC curves obtained for the two visual stimuli (left plot: for stimulating left visual hemifield; right plot: for stimulating right visual hemifield) when LG-RBSN were used (blue curve) versus ANTS (red curve). Our LG-RBSN method shows an AUC equal to 0.7053/0.7099 for left/right hemifield visual stimulation which demonstrates about 9.14%/7.02% improvement in comparison to the AUC obtained from ANTS ROC (0.6462/0.6634 for left/right visual hemifield stimulation). This result indicates that such improvement in the sensitivity of our proposed method is not at the expense of an increased false-positive rate.

**Fig. 12.**
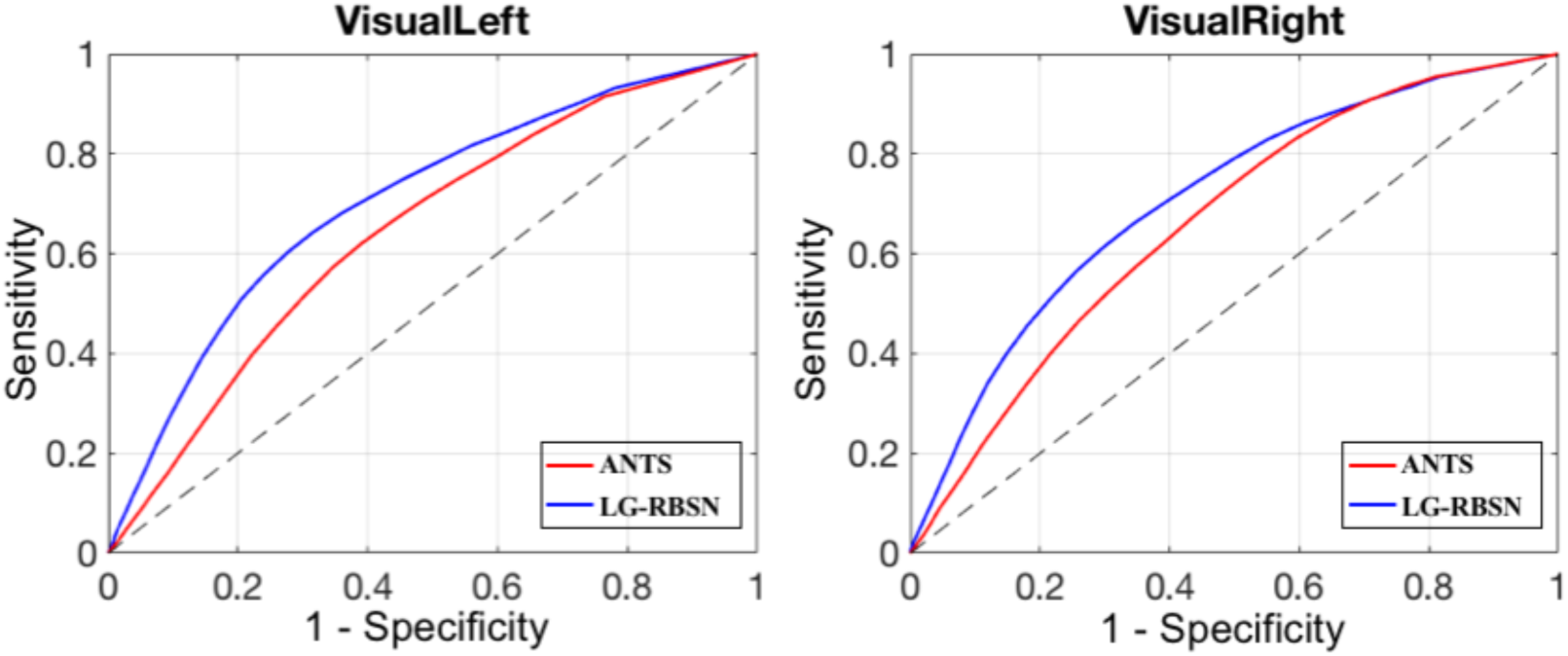
ROC curve evaluating spatial normalization methods with visual task-based fMRI group level t-statistics activation map compared to FreeSurfer lateral-occipital region (Left: for stimulating left visual hemifield; Right: for stimulating right visual hemifield).

Figure 13 illustrates the ROC curves obtained for the two auditory stimuli (left plot: for stimulating right hemisphere; right plot: for stimulating left hemisphere) when LG-RBSN were used (blue curve) versus ANTS (red curve). Our LG-RBSN method shows an AUC equal to 0.8234/0.8465 for left/right ear auditory stimulation which demonstrates about 0.95%/4.80% improvement in comparison to the AUC obtained from ANTS ROC (0.8156/0.8078 for left/right ear auditory stimulation). This result again indicates that such improvement in the sensitivity of our proposed method is not accompanied by an increase in the false-positive rate.

**Fig. 13.**
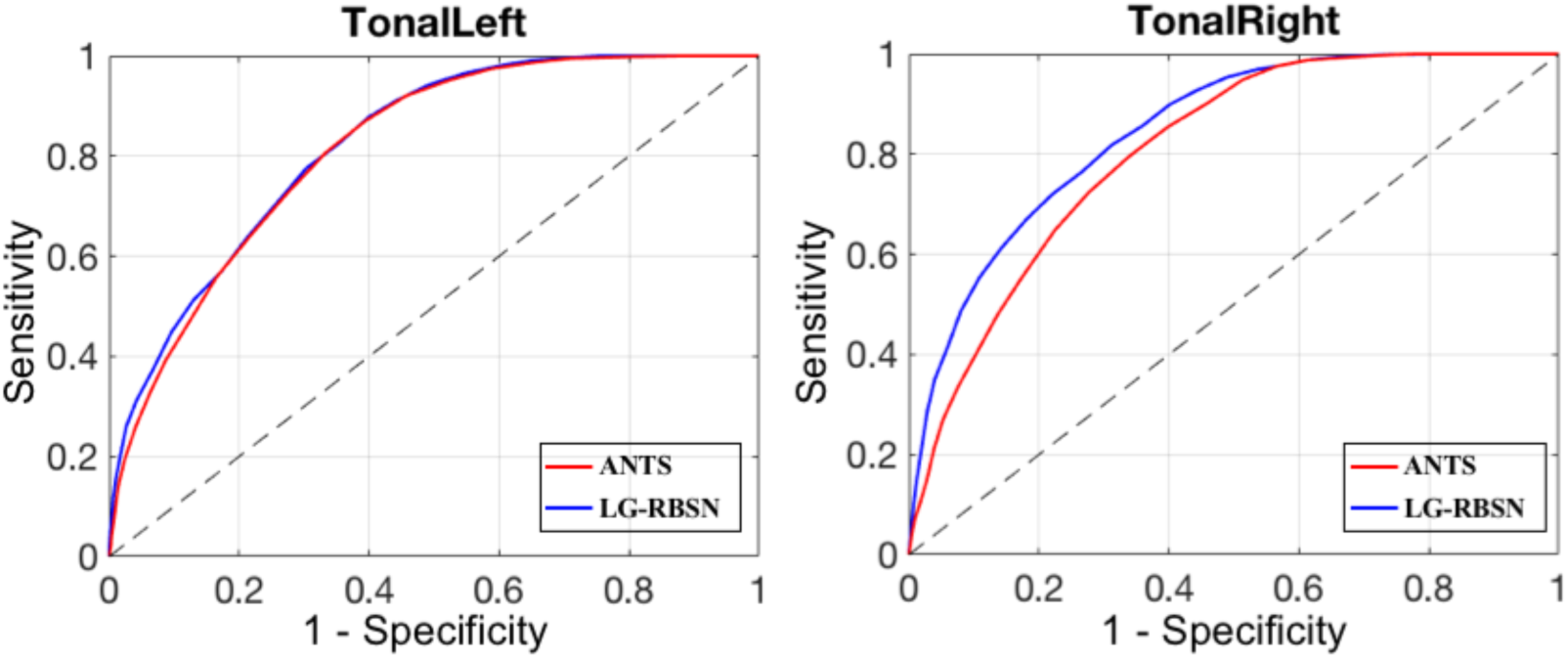
ROC curve evaluating spatial normalization methods with auditory task-fMRI group level T-statistics activation map compared to FreeSurfer segmented neuroanatomical brain contralateral transverse temporal mask (Left: for stimulating left ear; Right: for stimulating right ear).

Taken together, results in Figure 12 and Figure S.5 highlight the superiority of the proposed LG-RBSN method in detecting small effects that were not detectable using conventional methods such as ANTS, without increasing the false-positive rate, which is the ultimate goal in any advancement in method development for fMRI processing pipeline.

### D. Coding and Execution Time

Most of the implementation in this work is done with Matlab R2017b. We have shared the code for LG-RBSN method on our lab’s github repository (https://github.com/QuantitativeNeuroimagingLaboratory). The current implementation of the LG-RBSN takes an average of around 18 hours to be performed on each region. However, since the registration of each region is done independently, they all can be done in parallel using cluster of high-performance computing with around 100 number of cores. Using the cluster makes the total execution time to be around 75 hours for each subject’s brain registration. However, applying the obtained warping field takes no more time than applying other conventional spatial normalization methods.

## IV. DISCUSSION

We presented a novel volumetric spatial normalization method for human structural and functional brain image processing and statistical analysis. Compared to typical volume-based spatial normalization methods that use intensity-based similarity measurements, our method uses landmark guidance which specifies a concrete spatial correspondence in the volumetric space based on matching of the brain cortical folding patterns. To address the local minimum problem in optimization steps in most of the whole brain volume- or surface-based, or hybrid image registrations, we propose to independently estimate a diffeomorphic warping field for each cortical region. Combining the regional warping fields using the IDW interpolation technique generated a smooth global warping field that resulted in a smooth transition across adjacent regions. To impose bijectivity and topology preserving properties onto the global warping field, we proposed a forward and reverse warping fields residual compensation method, which is applicable to any other existing non-topology preserving deformation fields, as long as both forward and backward warpings are available.

We have evaluated our proposed method using three different experiments: 1) Using simulated 2D images of a single gyrus we demonstrated that our method not only aligns cortical folding patterns, but also keeps an accurate correspondence to its internal regional structures. Furthermore, we showed that our method is more robust to the local minimum problem compared to a top performing volumetric registration method. 2) Using structural images of human brains, we showed that our method increases the correspondence between cortical regions, sub-cortical regions, and cerebral white matter in comparison to the existing top performing algorithm. Our method also showed that the number of non-positive Jacobian voxels decreases with residual compensation iterations. 3) Using functional images of human brains, we first showed that improving the correspondence between regional structure not only increases the statistical power in detecting smaller activations, but also keeps the false-positive rate low. This was measured by AUC of the ROC curves, indicating about 8% improvement. Together, our findings suggest that LG-RBSN method is a more accurate and reliable substitute for conventional spatial normalization techniques commonly used in the field.

Compared to typical surface-based spatial normalization methods that only warp brain cortical surface, our method extended surface-based methods to estimate a volumetric warping field. This method is considered more robust than typical surface-based spatial normalization methods, since fMRI brain activations are captured originally in the 3D Euclidean space and thus avoiding the projection of the volumetric data onto the cortical surface. Furthermore, surface-based methods are more susceptible to the inaccuracies that often occur during reconstructions of cortical surface due to geometric distortion, especially in high-field and multiband acquisitions. While any inaccuracy in the surface reconstruction will cause surface-based method to include regions from outside of the brain or the white matter, our method computes volumetric deformation mappings that are applied to all gray-matter, white-matter, and other brain regions, thus preventing loss of fMRI data due to inaccurate surface extraction.

Evaluating spatial normalization methods has received considerable attention in recent years, which are typically evaluated by measuring the overlap between the warped anatomical regions and the counterpart regions in the reference brain. Klein *et al*. evaluated 14 methods with four MRI dataset of healthy and young subjects, and showed that these methods perform well in warping the sub-cortical regions (average DSC above 80%), but even the top performing method ANTS [5] generally has poor performance in warping cortical regions (average DSC between 60% and 70%) [13]. The recently-developed deep learning methods still have comparable performance as ANTS [6], [7]. It is because almost all of the widely used brain image registration techniques that work in the 3D Euclidean space, whether volume-based, or surface and volume hybrid methods, are based on solving the optimization problem of matching the whole brain at once and suffer from the local minimum problem, resulting in poor registration of brain cortical regions. Spatial normalization is even more challenging to studies of populations with severe brain morphology changes. For example, caution should be taken when studying aging population, as it has been shown that brain morphology changes along normal aging with grey matter volume reduction [35], especially in prefrontal regions [36] and in the medial temporal lobe [37]. In these cases, inaccurate spatial normalization can transfer population-related residuals to the normalized group-level brain activation, which will no longer be a valid representation of the population. Hence, any conclusion drawn with this biased representation will heavily be confounded by the residuals. For example, the age-related atrophy in the brain of the older participants have shown to further deteriorate the accuracy of the spatial normalization, and subsequently interpreted as age-related attenuation of BOLD response amplitude [38].

We have preliminarily used region-based registration methods to align MRI brain images with optimization running separately for each individual brain region instead of the whole brain [16]. This region-based spatial normalization method resulted in a 44% improvement of the correspondence between cortical regions (DSC around 0.75) in comparison to the state-of-the-art non-linear whole brain registration (ANTS). However, the inter-subject variability of cortical regions was still causing the optimization process to fall into local minimums especially in regions with sever age-related atrophy and deformations, like temporal lobe regions. Additionally, separately warping each region individually with its own warping field can introduce gaps and overlaps in-between regions. Edwards et al. has also compartmentalized the medical images of the human body into two separate segments in which the rigid and deformable structures of the body were registered independently to their counterparts, transformed separately, and then combined [39]. This method improved registration accuracy but was limited for the resultant discontinuity in structural boundaries. This limitation was addressed in a following paper [40] using a single global smooth transformation. The global transformation was composited using a modified radial basis function and the inverse distance interpolation [25] based on rigid structures within the image. Another similar method used only affine transformations for different segments of the brain [29]. In this work, each gyrus locally registered to its counterpart using affine transformation for 2D registration of myelin-stained histological sections of the human brain. Local registration methods interpolate locally linear transformation fields, whereas our method not only uses topographical landmarks to guide the non-linear registration, but also deals with the regional transition by an IDW interpolation method with enforced bijectivity to ensure the final global warping field is a diffeomorphic and topology preserving deformation.

The tradeoff between regional correspondence and bijectivity regularization (for topology preserving transformation) can be tuned to find a balance between the matching of brain structures and number of voxels violating the diffeomorphism. In our method, the number of non-positive Jacobian voxels can be lowered even to reach zeros, however, this will cause a decrease in the correspondence between the cortical regions (the average DSC of cortical region will drop to around 80%). In our experiments, we choose to tolerate 1300 number of voxels (0.0077% of total number of voxels) with non-positive Jacobian determinant, to achieve about 86% correspondence between cortical regions, which was the optimal setting in this work, however other applications may require further adjustment to obtain their optimal ranges. In addition, in the IDW interpolation method we set *μ* = 4, which produced acceptable results in our experiments. However, a thorough optimization is required in the future to obtain the optimal value for the *μ* in each registration application.

The reported DSC for existing methods in this work may seem lower than the ones reported in the literature. This is because; a) healthy elderly adults comprise more than one-third of our sample and generally show excessive brain atrophy in comparison to younger subjects, particularly in the prefrontal and temporal cortical regions [36], [37]. It has been shown that brain atrophy can significantly alter the effectiveness of brain registration accuracy [5]. b) we have evaluated a subject-to-template registration while most of the evaluation in the existing evaluation are done with subject-to-subject registration [13]. This will be even more problematic when we have older population in our subjects group. In future studies, cerebral cortex regions overlap can be further improved with a custom group average template specifically designed for use with LG-RBSN.

It is already known that the functional architecture of the brain does not necessarily and accurately follow brain gyri and sulci morphology. Therefore, one might conclude that improving the correspondence between brain structural features (sulci and gyri) might not necessarily translate to improvement in functional correspondence of the aligned regions, and would alter the effects of the improved spatial normalization methods in functional imaging of the brain. Since we currently do not have an accurate measurement of the deviation of the functional architecture from brain morphology, it is difficult to assess any limit in which improvement of the regional correspondence becomes unattainable. Nonetheless, we have shown that increasing the regional correspondence to 86% still increases the functional correspondence, and indicates that we are still operating under such limitation, if exists. Additionally, the proposed LG-RBSN can also be modified to identify landmarks from functional or other imaging modality data, so that landmarks correspondence is established using functional cortical registration methods to help align the brain functional organizations across subjects.

In future work, it would be interesting to evaluate the performance of the LG-RBSN method for multivariate techniques such as group independent component analysis (ICA), or partial least-squares (PLS). The difference between the evaluation of the multivariate and univariate (voxel-based) methods is that multivariate techniques often require warping the actual 4D fMRI data, whereas univariate analysis can be performed after the first-level statistical analysis. Therefore, one might expect to obtain a different performance in applying LG-RBSN to the multivariate data. Furthermore, we only evaluated the LG-RBSN technique on structural and functional MRI scans. It would be interesting to assess the effectiveness of this method on other MRI modalities such as diffusion weighted imaging, arterial spine labeling, and susceptibility weighted imaging amongst others. We also expect that our proposed spatial normalization method could be extended to enhance spatial normalization accuracy in the other imaging modalities such as positron emission tomography and computed tomography.

## V. CONCLUSION

Using automatic landmark detection and matching, we have developed and implemented a novel 3D volumetric spatial normalization method that not only aligns the cortical folding patterns of the brain, but also results in a high correspondence between different regions along the cortical ribbon and their counterparts in the template image. Our method substantially outperforms the existing top performing spatial normalization method by giving a significantly higher correspondence between the structure of the neuroanatomical regions, and also yields higher sensitivity and specificity in the group-level statistics when analyzing task-based fMRI data with both auditory and visual stimuli. The limited accuracy of conventional methods becomes more prominent when applied to clinical and aging populations with severe alterations in brain morphology. When compared to the healthy group, this population-based bias has been shown to generate false-positive findings that have been reported as a genuine breakthrough in the literature. We conclude that our proposed LG-RBSN method is a suitable substitute for the conventional volumetric whole brain registration methods that often fail to generate an accurate correspondence between regions of the cerebral cortex, particularly for clinical and aging populations.

## Acknowledgment

The authors would like to thank Emily B. Tanzi for paper proofreading.

## Notes

### Competing Interest Statement

The authors have declared no competing interest.

